# Effects of 7.83 Hz and 528 Hz Extremely Low-Frequency Electromagnetic Fields on Early Growth of Turnip Greens *Brassica rapa* var. *rapa*

**DOI:** 10.1101/2025.11.11.687793

**Authors:** Igor Nelson

## Abstract

Extremely low-frequency (ELF) electromagnetic fields are known to interact with biological systems, but their effects on plant development remain incompletely characterized. This study examined the influence of weak ELF electromagnetic fields on the early growth of *Brassica rapa* var. *rapa* (turnip greens). Seeds were germinated under controlled environmental conditions and exposed for 60 minutes per day over five consecutive days to alternating electromagnetic fields at 7.83 Hz or 528 Hz, generated by solenoidal coils. Germination rates were not significantly affected by either treatment. However, seedlings exposed to both frequencies exhibited significantly greater shoot elongation compared with unexposed controls. These findings indicate that weak ELF electromagnetic fields can promote early vegetative development in *B. rapa* var. *rapa*. without altering germination success. Further studies involving multiple species, improved field characterization, and molecular assays are warranted to clarify the mechanisms underlying these effects.

## 1. Introduction

Biological systems have evolved in constant interaction with natural electromagnetic fields, including solar radiation, the geomagnetic field, and the Earth’s extremely low-frequency (ELF) oscillations (Nelson 2025). These fields are persistent environmental factors that have been present throughout the evolution of life and are known to influence various physiological processes. The Earth’s ionosphere cavity supports the Schumann resonance spectrum (fundamental mode around 7.83 Hz and higher harmonics), a naturally occurring ELF background generated by global lightning activity, which has been discussed as a potential synchronizing environmental factor for biological rhythms (Nelson 2025; Cherry 2002; Volland 1995).

Several reviews report that weak ELF exposures can modulate ion transport, membrane potential, and intracellular Ca^2+^ signaling in various cell types, though these effects are generally small and often dependent on exposure parameters and biological context (Pall 2013; Panagopoulos et al. 2015; Tian 2023). Despite decades of research, the precise biophysical mechanisms remain incompletely understood.

Experimental studies have reported modest biological effects at or near Schumann’s main mode (7.83 Hz), such as modulation of intracellular calcium and changes in human EEG coherence. In addition, several in vitro studies have suggested potential effects on cancer cell proliferation and signaling: for example, exposure to this frequency been shown to influence the growth of human breast cancer cells (Wang et al. 2021), and 7.83 Hz square-wave fields can inhibit B16F10 melanoma cell growth via L- and T-type calcium channels (Wang et al. 2020; Tang et al. 2019). However, results remain preliminary and not always reproducible.

Since plants are stationary and highly sensitive to environmental conditions, they serve as excellent models for studying non-thermal effects of electromagnetic fields (Vian et al. 2016).

Accumulating evidence points in the direction that nonionizing electromagnetic fields can affect plant germination, growth, photosynthetic performance, and secondary metabolism (Vian et al. 2016; Galland and Pazur 2018; Roux et al. 2006). However, the amplitude and reproducibility of these effects vary considerably depending on field strength, frequency, and plant species (Vian et al. 2016; Maffei 2014).

Other than 7.83 Hz, the frequency 528 Hz is typically discussed in the context of acoustic (sound) rather than electromagnetic stimulation. Although popularized as a “healing frequency,” empirical data are limited. One controlled human study found that listening to music tuned to 528 Hz reduced salivary cortisol and increased oxytocin compared with 440 Hz music (Akimoto 2018). Another experiment using cultured human astrocytes reported increased cell viability and decreased oxidative stress following 528 Hz sound exposure (Babayi and Riazi 2017). In rodents, prolonged 528 Hz auditory exposure is known to be associated with reduced anxiety-like behaviors and altered neuroendocrine markers (Babayi Daylari et al. 2019).

These findings pertain to acoustic bioeffects, not to ELF electromagnetic interactions, but motivate broader exploration of frequency-specific biological effects.

Several mechanisms have been proposed to explain how weak ELF fields might influence biological systems, including modulation of voltage-gated ion channels, resonance with calciumbinding kinetics, and effects on reactive oxygen species and redox signaling (Pall 2013; Binhi and Rubin 2020). Although none are conclusively demonstrated, such hypotheses guide alternative experimental designs.

In this context, the present study investigates the influence of extremely low-frequency electromagnetic field at 7.83 Hz and 528 Hz on seed germination, shoot growth in *Brassica rapa* var. *rapa*. The selected frequencies (7.83 Hz and 528 Hz) represent, respectively, a natural ELF background resonance and a common “healing” frequency with reported psychophysiological effects.

## 2. Materials and Methods

### 2.1. Overview

Turnip greens (*B. rapa* var. *rapa*) seeds were germinated under controlled environmental conditions. Five groups (n = 30 per group) were maintained under identical conditions of natural light, temperature, and humidity throughout the experiment. Seeds were germinated on cotton disks placed in plastic cups, each sample initially watered with 4 mL of water on the first day and 1 mL of water per day on the following 4 days.

The experimental groups were exposed to alternating electro-magnetic fields (AC) for 60 minutes per day over five consecutive days at 7.83 Hz and 528 Hz EMF. All samples were kept on the same shelf throughout the experiment; during exposure periods, the experimental groups were positioned on their respective active coils, while the control group remained on the shelf approximately 4 m away from the coils. Frequencies of 7.83 Hz and 528 Hz were selected due to prior reports of biological relevance.

Samples were rotated within the same positions throughout the experiment, ensuring that each group passed through all positions to minimize positional bias. After five days, shoot length (cm) was measured using a millimetric ruler with an estimated precision of *±*1 mm, accounting for curvature and handling variability. Mean shoot length per group was calculated from germinated seedlings for statistical comparison.

### 2.2. Coil Configuration

Two continuous alternating current sinusoidal signals were generated using Audacity software (Audacity Team, open-source audio editor), respectively at 7.83 Hz and 528 Hz. Each frequency was supplied independently to a dedicated TDA2050-based audio amplifier. The amplifiers were connected through 3.5 mm audio jack outputs from a computer and each powered by its own power supply.

#### Electromagnetic Setup

The field source was a solenoidal-type coil wound with 0.20 mm enamelled copper wire, with an approximate total wire length of 100 m. The coil was wound around a circular former with an inner diameter of 5.2 cm, an outer diameter of 6.5 cm, and a coil height of 0.9 cm. The measured DC resistance of the coil was approximately 79 Ω. From the measured resistance and the resistivity of copper at 20^*°*^C, the estimated number of turns was approximately *N* ≈790.

The coil was powered by a regulated DC supply rated at 2 V, 4.5 A and driven with a sinusoidal signal amplified by a TDA2050-based power amplifier. Two excitation frequencies were applied: 7.83 Hz and 528 Hz.

The voltage and current delivered to the coil were measured using a true RMS ammeter and an oscilloscope. At 7.83 Hz, the applied signal measured 14.3 V_pp_ and 0.317 A, while at 528 Hz, the measurements were 10.10 V_pp_ and 2.37 A, respectively.

The magnetic field was measured with a MAG3110 sensor at 7.83 Hz and was found to be approximately 1 mT at the location of the sample. Note that the sensor is not capable of measuring the magnetic field at 528 Hz.

The electric field was measured using a copper plate connected to an Arduino and was estimated to be approximately 33 V*/*m at 7.83 Hz and 29 V*/*m at 528 Hz at the sample location.

### 2.3. Results and Analysis

#### 2.3.1. Germination Analysis

For all groups, a total of 30 seeds were planted.

As shown in Table 1, germination outcomes were analyzed separately from shoot-length measurements. A chi-square test of independence was performed on the 3 *×* 2 contingency table (germinated vs. non-germinated across groups), showing no significant difference in germination rates among treatments (*χ*_2_(2) = 2.61, *p* = 0.271). All expected cell counts ( 7) exceeded the conventional threshold of 5, confirming the validity of the chi-square approximation.

**Table 1.**
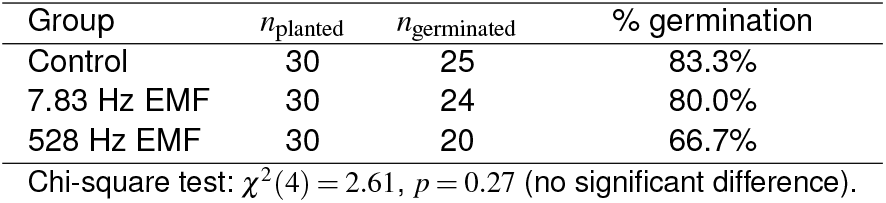
Germination counts and chi-square test (assumed *n*_planted_ = 30 per group).

These results indicate that exposure to electromagnetic fields did not significantly affect seed germination rates under the tested conditions, suggesting that any subsequent differences observed in shoot growth are unlikely to result from differential germination success.

Data were analyzed using independent-samples *t*-tests (with unequal variances not assumed). Analyses were restricted to germinated seeds (sample sizes as shown in Table 2). Normality was evaluated using the Shapiro–Wilk test. All experimental groups except the control showed no significant deviation from normality (*p >* 0.05). Because the control group showed a departure from normality (Shapiro–Wilk *p* = 0.008), Welch’s two-sample *t*-tests were used (robust to unequal variances and sample sizes) to compare mean shoot lengths, and confirmed all significant results using nonparametric Mann–Whitney *U* tests. Means ± SD, normality *p*-values, and both parametric and nonparametric comparisons against the control group are reported.

**Table 2.**
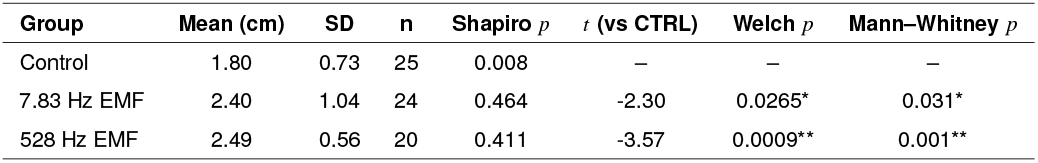
Descriptive and inferential statistics for shoot length across treatments.

| Group | Mean (cm) | SD | n | Shapiro $p$ | $t$ (vs CTRL) | Welch $p$ | Mann–Whitney $p$ |
| --- | --- | --- | --- | --- | --- | --- | --- |
| Control | 1.80 | 0.73 | 25 | 0.008 | – | – | – |
| 7.83 Hz EMF | 2.40 | 1.04 | 24 | 0.464 | -2.30 | 0.0265* | 0.031* |
| 528 Hz EMF | 2.49 | 0.56 | 20 | 0.411 | -3.57 | 0.0009** | 0.001** |

Field-exposed groups exhibited significantly greater shoot elongation than the unexposed control (*p <* 0.05).

**Figure 1.**
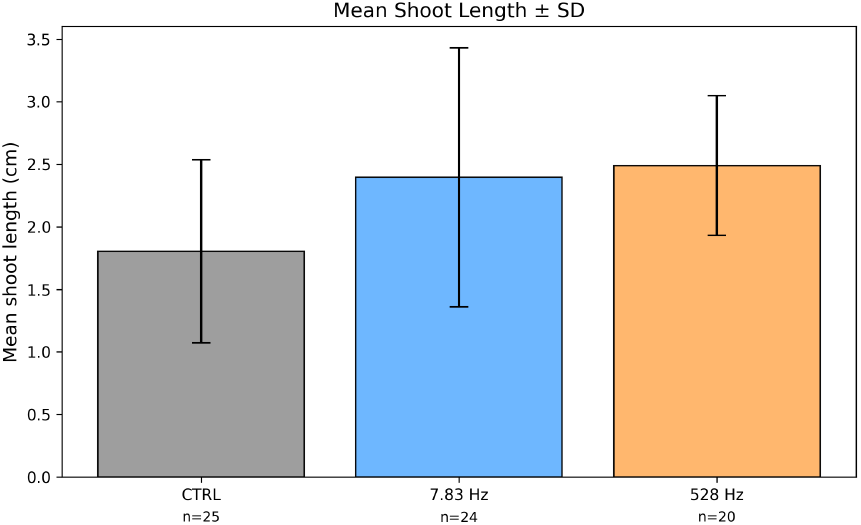
Mean shoot length (cm) for each experimental group. Error bars represent standard deviations. Sample sizes are shown above each bar (*n* = number of germinated seedlings). Groups correspond to: Control, 7.83 Hz and 528 Hz.

## 3. Discussion

This study investigated the effects of weak ELF electromagnetic fields on early growth of *B. rapa* var. *rapa*. Results demonstrate that while germination percentages were not significantly affected by any treatment, all field-exposed groups exhibited enhanced shoot elongation relative to the unexposed control. This finding supports the idea that weak ELF electromagnetic fields can modulate early vegetative growth under controlled environmental conditions.

Given the low power levels and negligible temperature increase of the applied fields, the observed differences are unlikely to be thermal in origin. Potential explanations include modulation of ion transport, changes in membrane polarization, or resonance-like interactions near the 7.83 Hz Schumann frequency, consistent with previous studies reporting biological responses to weak ELF fields Nelson (2025); Pall (2013); Binhi and Rubin (2020). Further investigations with higher replication, individual plant imaging, and additional physiological measurements (e.g., chlorophyll content, root growth, or gene expression) are needed to clarify the mechanisms and confirm the preliminary observations reported here.

Overall, the present findings suggest that weak ELF exposures can enhance early vegetative development in *B. rapa* var. *rapa*.

While the biological significance and reproducibility of these results require further study, they provide a foundation for future research exploring the influence of weak electromagnetic fields on plant development.

The exposure magnitudes and durations in this study were consistent with or lower than those used in prior plant research. For example, previous studies investigating ELF effects on seed germination and early growth often applied magnetic fields ranging from 0.1 to 5 mT for 30–120 minutes per day (Vian et al. 2016; Sarraf et al. 2020). In our experiments, the solenoidal coil generated approximately 1 mT at 7.83 Hz. Exposure durations of 60 minutes per day over five consecutive days fall within the lower-to-mid range of previous protocols, suggesting that the observed growth effects occurred at relatively modest field intensities and durations.

Several limitations of the current study should be acknowledged. The experiment involved a single plant species, limiting generalizability to other species. Sample sizes were modest, and no blinding was applied during measurements, introducing potential bias. Additionally, although care was taken to position control samples away from active coils, ambient electromagnetic fields in the laboratory could have contributed to background exposure.

## 4. Conclusions

The aim of this study was to investigate the effects of different electromagnetic frequencies on the early growth of *B. rapa* var. *rapa*.

Germination percentage was not affected by any of the treatments.

All examined field-exposed groups showed increased shoot elongation relative to the control group, supporting the hypothesis that weak ELF electromagnetic fields can enhance early plant growth under controlled environmental conditions.

Due to the low power levels and negligible temperature increase of the applied fields, the observed differences are unlikely to be thermal in origin.

The solenoid coil generated a magnetic field of approximately 1 mT and an electric field of 33 V/m at 7.83 Hz.

Further studies with higher replication and more detailed imaging are needed to confirm these preliminary observations and clarify the underlying mechanisms.

## 5. Future Work

To build on the findings reported here, future studies should incorporate electromagnetic shielding and real-time field mapping to isolate the contribution of coil geometry more rigorously. Increasing replication and including multiple plant species will help assess the generalization of ELF and other field effects. Additional physiological and molecular endpoints such as chlorophyll content, root development, or gene expression, should be measured to clarify the mechanisms underlying growth modulation. Longer-term experiments may also determine whether early vegetative development enhancements translate into differences in mature plant development or yield.

## Declaration of generative AI and AI-assisted technologies in the manuscript preparation process

During the preparation of this work the author used ChatGPT (GPT-5, OpenAI, 2025) in order to assist with background research and to improve English writing and readability. After using this tool/service, the author reviewed and edited the content as needed and takes full responsibility for the content of the published article.

## References

Akimoto, T. e. a. (2018). Effects of 528 hz music on the endocrine system and autonomic function. Health, 10(3):115–122.

Babayi, T. and Riazi, G. H. (2017). The effects of 528 hz sound wave to reduce cell death in human astrocyte primary cell culture treated with ethanol. Journal of Addiction Research and Therapy, 2017:1–5.

Babayi Daylari, T., Riazi, G. H., Pooyan, S., Fathi, E., and Hedayati Katouli, F. (2019). Influence of various intensities of 528 hz sound-wave in production of testosterone in rat’s brain and analysis of behavioral changes. Genes & Genomics, 41(2):201–211.

Binhi, V. N. and Rubin, A. B. (2020). Molecular gyroscopes, biological water, and quantum effects of weak magnetic fields. Biophysical Reviews, 12:1031–1041.

Cherry, N. J. (2002). Schumann resonances, a plausible biophysical mechanism for the human health effects of solar and geomagnetic activity. Natural Hazards, 26:279–331.

Galland, P. and Pazur, A. (2018). Magnetic field effects on plant growth, development, and evolution. Frontiers in Plant Science, 9:181.

Maffei, M. E. (2014). Magnetic field effects on plant growth, development, and evolution. Frontiers in Plant Science, 5:445.

Nelson, I. (2025). Exploring the influence of schumann resonance and electromagnetic fields on bioelectricity and human health. Electromagnetic biology and medicine, 44(3):348–358. doi:10.1080/15368378.2025.2508466.

Pall, M. L. (2013). Electromagnetic fields act via activation of voltage-gated calcium channels to produce beneficial or adverse effects. Journal of Cellular and Molecular Medicine, 17(8):958–965.

Panagopoulos, D. J., Johansson, O., and Carlo, G. L. (2015). Mechanism for action of electromagnetic fields on cells. Biochemical and Biophysical Research Communications, 460:97–102.

Roux, D., Vian, A., Girard, S., Bonnet, P., Paladian, F., Davies, E., and Ledoigt, G. (2006). Electromagnetic fields (900 mhz) evoke consistent molecular responses in tomato plants. Physiologia Plantarum, 128:283–288.

Sarraf, M., Kataria, S., Taimourya, H., Santos, L. O., Menegatti, R. D., Jain, M., Ihtisham, M., and Liu, S. (2020). Magnetic field (mf) applications in plants: An overview. Plants, 9(9):1139.

Tang, J. Y., Yeh, T. W., Huang, Y. T., Wang, M. H., and Jang, L. S. (2019). Effects of extremely low-frequency electromagnetic fields on b16f10 cancer cells. Electromagnetic Biology and Medicine, 38(2):149–157.

Tian, F. e. a. (2023). Extremely low-frequency magnetic fields modulate intracellular calcium signaling: A review. Environmental Research, 230:115401.

Vian, A., Davies, E., Gendraud, M., and Bonnet, P. (2016). Plant responses to high frequency electromagnetic fields. BioMed Research International, 2016:1830262. FEMU ID: 29052; EMF-Portal URL: https://www.emfportal.org/en/article/29052.

Volland, H., editor (1995). Handbook of Atmospheric Electrodynamics, Volume I. CRC Press, Boca Raton, FL, 1st edition.

Wang, M. H., Chen, K. W., Ni, D. X., Fang, H. J., Jang, L. S., and Chen, C. H. (2021). Effect of extremely low frequency electromagnetic field parameters on the proliferation of human breast cancer. Electromagnetic Biology and Medicine, 40(3):384–392.

Wang, M. H., Jian, M. W., Tai, Y. H., Jang, L. S., and Chen, C. H. (2020). Inhibition of b16f10 cancer cell growth by exposure to the square wave with 7.83 ± 0.3 hz involves l- and t-type calcium channels. Electromagnetic Biology and Medicine, 40(1):150–157.

